# Broadly Reactive H2 Hemagglutinin Vaccines Elicit Cross-Reactive Antibodies in Ferrets Pre-Immune to Seasonal Influenza A Viruses

**DOI:** 10.1101/2021.02.17.431747

**Authors:** Z. Beau Reneer, Amanda S. Skarlupka, Parker J. Jamieson, Ted M. Ross

## Abstract

Influenza vaccines have traditionally been tested in naïve mice and ferrets. However, humans are first exposed to influenza viruses within the first few years of their lives. Therefore, there is a pressing need to test influenza virus vaccines in animal models that have been previously exposed to influenza viruses before being vaccinated. In this manuscript, previously described H2 computationally optimized broadly reactive antigen (COBRA) HA vaccines (Z1, Z5) were tested in influenza virus ‘pre-immune’ ferret models. Ferrets were infected with historical, seasonal influenza viruses to establish pre-immunity. These pre-immune ferrets were then vaccinated with either COBRA H2 HA recombinant proteins or WT H2 HA recombinant proteins in a prime-boost regimen. A set of naïve pre-immune or non pre-immune ferrets were also vaccinated to control of the effects of the multiple different pre-immunities. All of the ferrets were then challenged with a swine H2N3 influenza virus. Ferrets with pre-existing immune responses influenced recombinant H2 HA elicited antibodies following vaccination as measured by HAI and classical neutralization assays. Having both H3N2 and H1N1 immunological memory regardless of the order of exposure significantly decreased viral nasal wash titers and completely protected all ferrets from both morbidity and mortality, including the mock vaccinated ferrets in the group. While the vast majority of the pre-immune ferrets were protected from both morbidity and mortality across all of the different pre-immunities, the Z1 COBRA HA vaccinated ferrets had significantly higher antibody titers and recognized the highest number H2 influenza viruses in a classical neutralization assay compared to the other H2 HA vaccines.

**Importance:** H1N1 and H3N2 influenza viruses have co-circulated in the human population since 1977. Nearly every human alive today has antibodies and memory B and T cells against these two subtypes of influenza viruses. H2N2 influenza viruses caused the 1957 global pandemic and people born after 1968 have never been exposed to H2 influenza viruses. It is quite likely that a future H2 influenza virus could transmit within the human population and start a new global pandemic, since the majority of people alive today are immunologically naïve to viruses of this subtype. Therefore, an effective vaccine for H2 influenza viruses should be tested in an animal model with previous exposure to influenza viruses that have circulated in humans. Ferrets were infected with historical influenza A viruses to more accurately mimic the immune responses in people who have pre-existing immune responses to seasonal influenza viruses. In this study, pre-immune ferrets were vaccinated with WT and COBRA H2 recombinant HA proteins in order to examine the effects of pre-existing immunity to seasonal human influenza viruses have on the elicitation of broadly cross-reactive antibodies from heterologous vaccination.

## Introduction

The 1957 ‘Asian Influenza’ pandemic was caused by an H2N2 influenza virus resulting in an estimated one to two million deaths worldwide (1). This novel H2N2 influenza virus was the result of a reassortment event between a human H1N1 influenza virus and an avian H2N2 influenza virus (2). This novel H2N2 influenza virus contained the HA, NA, and PB1 genome segments from an avian H2N2 influenza virus and the other five genome segments from a human H1N1 influenza virus (3). The 1889 influenza pandemic may also have been caused by an H2N2 influenza virus (4). Therefore, at least one of the last five influenza pandemics was caused by an influenza virus from the H2N2 subtype, it is likely that a future pandemic will be caused by an H2N2 influenza virus.

H2 influenza viruses have not been as extensively studied as other influenza A virus subtypes such as H1, H3, H5 or H7. While H2 influenza viruses have been isolated numerous times from wild avian species and domestic poultry (5–10), there have been no known viral infections of humans since the 1960s. However, in 2006 a novel H2 influenza virus was isolated from two separate swine farms in Missouri (11). This swine derived H2N3 influenza virus has been shown to cause severe disease in both mice and ferrets (12). The H2 HA is also capable of obtaining a multi-basic cleavage site and remaining functional which could have dire implications for both humans and poultry in the future (13).

The goal of this study was to evaluate how memory immune responses to previous influenza virus infections affect broadly-reactive HA-based vaccinations. To develop broadly reactive influenza virus vaccines, our group has used the methodology for enhanced antigen design, termed computationally optimized broadly reactive antigen (COBRA) to design hemagglutinin (HA) immunogens for the H1, H3, and H5 influenza subtypes (14–21). This process utilizes multiple rounds of layered consensus building to generate influenza virus vaccine HA antigens that are capable of eliciting broadly reactive HA antibodies that can protect against both seasonal and pandemic influenza strains that have undergone genetic drift (17, 18, 21). These vaccine antigens also inhibit viral infection and virus induced pathogenesis in mice, ferrets, and non-human primates (16, 22–24). Using the consensus layering approach of COBRA design, H2 COBRA HA vaccines were previously developed and characterized (25).

Humans have been infected with different types of influenza viruses throughout their lives (3). Additionally, different subtypes have circulated as seasonal influenza viruses over the past 100 years (3). A person’s history of influenza virus infections has an effect on future influenza vaccinations and infections (26–28). The HA subtype from the first influenza virus infection influences the susceptibility of an individual to subsequent influenza virus infections from other subtypes (27). Therefore, a broadly cross-reactive H2 influenza virus vaccine should be evaluated in animal previously infected or ‘pre-immune’ to different influenza virus subtypes.

For this study, fitch ferrets were infected with different combinations of human isolated H1N1 and H3N2 influenza viruses. These two influenza virus subtypes are the only influenza A viruses that have circulated in the human population since 1968 and would therefore be reflective of the majority of individuals alive today. The H1N1 infections included both a seasonal (before 2009) and a pandemic H1N1 virus (2009-present) since individuals alive today who are over the age of eleven would have been exposed to both types of H1N1 influenza viruses. The H1N1 viruses used in this study were Singapore/6/1986 (Sing/86) and California/07/2009 (CA/09) respectively. The H3N2 influenza viruses used to establish pre-immunity were either Sichuan/2/1987 (Sich/87) or Panama/2007/1999 (Pan/99). Additionally, the influenza virus pre-immunity of individuals alive today would include individuals infected with H1N1 influenza viruses followed by H3N2 influenza viruses and vice-versa. Finally, a ‘non pre-immune’ or ‘naïve pre-immune’ group was included as a control for the vaccines alone. An H2N2 pre-immune group was also included as a pseudo “positive control” group since previous studies have shown that imprinting ferrets with a specific subtype of influenza virus followed by vaccination with another antigenically distinct influenza virus of the same subtype induces expansive intra-subtype antibodies (29). Two antigenically distinct H2N2 avian influenza viruses were used to establish the H2N2 pre-immunity because of the restrains of housing ferrets infected with BSL3 human H2N2 influenza viruses in high level containment for several months.

After pre-immunity was established, two H2 COBRA HA vaccines (Z1, Z5) were used to vaccinate the ferrets. Protective immune responses elicited by the Z1 and Z5 COBRA HA vaccines were compared to the elicited response in pre-immune ferrets vaccinated with wild-type H2 HA proteins. The Z1 COBRA HA vaccinated pre-immune ferrets showed more broadly cross-reactive antibody responses to a panel of H2 influenza viruses across each of the six pre-immune immune groups compared to ferrets vaccinated with either of the two wild-type H2 HA vaccines. Therefore, the Z1 COBRA HA would be an ideal vaccine for use in individuals regardless of their previous exposure to influenza A viruses.

## Materials and Methods

### Viruses, recombinant HA proteins and virus-like particles

A/Chicken/Potsdam/4705/1984 (Chk/Pots/84) (H2N2) (clade-1), A/Chicken/PA/298101-4/2004 (Chk/PA/04) (H2N2) (clade-1), A/Duck/Hong Kong/273/1978 (Duk/HK/78) (H2N2) (clade-2), A/Mallard/Minnesota/AI08-3437/2008 (Mal/MN/08) (H2N3) (clade-3), A/Swine/Missouri/4296424/2006 (Sw/MO/06) (H2N3) (clade-3), A/Formosa/313/1957 (For/57) (H2N2) (clade-2), and A/Taiwan/1/1964 (T/64) (H2N2) (clade-2) were obtained from either the United States Department of Agriculture (USDA) Diagnostic Virology Laboratory (DVL) in Ames, Iowa, from BEI resources (City, State, USA), or provided by the laboratory of Dr. S. Mark Tompkins (Athens, GA, USA). Each influenza virus was passaged using embryonated chicken eggs except for the Sw/MO/06 virus which was passaged in MDCK cells. Each influenza virus was harvested from either the eggs or cells and aliquoted into tubes which were stored at −80°C. Each influenza virus was tittered using a standard influenza plaque assay described below.

Recombinant HA (rHA) proteins were expressed using the pcDNA 3.1+ plasmid (Addgene, Watertown, MA). Each HA gene was truncated by removing the transmembrane (TM) domain and the cytoplasmic tail at the 3’ end of the gene (amino acids 527 to 562). The TM domain was determined using the TMHMM Server v. 2.0 website: http://www.cbs.dtu.dk/services/TMHMM/. The HA gene was truncated at the first amino acid prior to the TM domain. A fold-on domain from T4 bacteriophage (30), an Avitag (31) and a 6X histidine tag totaling 477 nucleotides were added to the 3’ end of the HA gene. The pcDNA 3.1+ vectors were then transfected individually into human embryonic kidney (HEK293T) suspension cells using ExpiFectamine 293 transfection reagent (ThermoFisher Scientific, Waltham, MA, USA) following manufacturer’s specifications. The supernatants (~500 mL) were then harvested from the transfected HEK293T cells. Each rHA was then purified from the supernatant using a nickel-agarose column (ThermoFisher Scientific, Waltham, MA, USA). The rHA proteins were then eluted from the column using 100mM imidazole (ThermoFisher Scientific, Waltham, MA, USA). After elution, the proteins were quantified using bicinchoninic assay (BCA) (ThermoFisher Scientific, Waltham, MA, USA) and stored at −80°C. Recombinant HA proteins produced for this study were A/Mallard/Netherlands/13/2001 (Mal/NL/01), Mallard/Wisconsin/08OS2844/2008 (Mal/WI/08), Z1 COBRA (Z1), and Z5 COBRA (Z5).

For the VLP production, adherent human endothelial kidney 293T (HEK-293T) cells were grown in complete Dulbecco’s modified eagles’ medium (DMEM) media. Once confluent, (1 × 10^6^) these cells were transiently transfected for the creation of mammalian virus-like particles (VLPs). Viral proteins were expressed from the pTR600 mammalian expression vectors (32). Influenza virus neuraminidase (A/South Carolina/1/1918; H1N1), the HIV p55 Gag, and HA expression plasmids expressing one of the H2 wild-type or H2 COBRA HA proteins were added to serum free media following the Lipofectamine 3000 protocol in a 1:2:1 ratio with a final DNA concentration of 1 ug. Following 72h of incubation at 37°C, supernatants from transiently transfected cells were collected, centrifuged to remove cellular debris, and filtered through a 0.22 μm pore membrane. VLPs were purified and sedimented by ultracentrifugation on a 20% glycerol cushion at 23,500 x g for 4h at 4°C. VLPs were resuspended in phosphate buffered saline (PBS), and total protein concentration was determined with the Micro BCA Protein Assay Reagent kit (Pierce Biotechnology, Rockford, IL, USA). Hemagglutination activity of each preparation of VLP was determined by serially diluting volumes of VLPs and adding equal volume 0.8% turkey red blood cells (RBCs) (Lampire Biologicals, Pipersville, PA, USA) suspended in PBS to a V-bottom 96-well plate with a 30 min incubation at room temperature (RT). Prepared RBCs were stored at 4°C and used within 72 h. The highest dilution of VLP with full agglutination of RBCs was considered the endpoint HA titer. The H2 HA sequences used for VLPs were Mal/NL/01, Chk/Pots/84, Muskrat/Russia/63/2014 (Musk/Rus/14) (clade-1), Duck/Cambodia/419W12M3/2013 (Duk/Cam/13) (clade-2), Japan/305/1957 (J/57) (clade-2), Moscow/1019/1965 (Mosc/65) (clade-2), T/64, Duk/HK/78, Mal/WI/08, Sw/MO/06, Quail/Rhode Island/16-018622-1/2016 (Qu/RI/16) (clade-3), Turkey/California/1797/2008 (Tk/CA/08) (clade-3).

### Ferret vaccination and challenge experiments

Fitch ferrets (*Mustela putorius furo*, spayed, female, 6 to 12 months of age), were purchased certified influenza free and de-scented from Triple F Farms (Sayre, PA, USA). Ferrets were pair housed in stainless steel cages (Shor-Line, Kansas City, KS) containing Sani-Chips laboratory animal bedding (P. J. Murphy Forest Products, Montville, NJ). Ferrets were provided with Teklad Global Ferret Diet (Harlan Teklad, Madison, WI) and fresh water *ad libitum*. The University of Georgia Institutional Animal Care and Use Committee approved all experiments, which were conducted in accordance with the National Research Council’s *Guide for the Care and Use of Laboratory Animals*, The Animal Welfare Act, and the CDC/NIH’s *Biosafety in Microbiological and Biomedical Laboratories* guide. Ferrets (*n = 20*) were pre-infected with H1N1, or H3N2 seasonal influenza viruses or H2N2 avian influenza viruses in different orders before vaccination. These influenza viruses included the H1N1 influenza viruses Singapore/6/1986 (Sing/86) and California/07/2009 (CA/09), the H3N2 influenza viruses Sichuan/2/1987 (Sich/87) or Panama/2007/1999 (Pan/99), and the H2N2 avian influenza viruses Chk/PA/04, or Qu/RI/16 all at an infectious dose of 1e+6 PFU in 1mL intranasally. For the ferrets with multiple pre-immune infections, ferrets were left for sixty days between each infection and before the first vaccination.

After the establishment of pre-immunity by viral infection, sixty days elapsed before ferrets were vaccinated with recombinant hemagglutinin (rHA) twice with four weeks between vaccinations. The ferrets were vaccinated with a 1:1 ratio (500μL total volume) of rHA diluted with phosphate-buffered saline (PBS) (15.0μg rHA/ferret) and the emulsified oil-water adjuvant, Addavax (InvivoGen, San Diego, CA, USA). The ‘mock’ vaccinated groups received only PBS and Addavax adjuvant at a 1:1 ratio (500μL total volume) with no rHA. Each vaccination was given intramuscularly. Before vaccinations and two weeks after each of the vaccinations, ferrets were bled and serum was isolated from each of the samples. The blood was harvested from all anesthetized ferrets via the anterior vena cava at days 0, 14, and 42. Blood samples were incubated at room temperature for 1h prior to centrifugation at 6,000 rpm for 10 minutes. The separated serum was removed and frozen at −20°C. The ferrets were infected four to six weeks after the second vaccination with the H2N3 influenza virus Swine/Missouri/4296424/2006 (Sw/MO/06). Animals were monitored daily for 10 days post-infection for clinical symptoms such as weight loss (20% n=3), lethargy (n=1), sneezing, dyspnea (n=2), and neurological symptoms (n=3). Any ferret that reached a cumulative score (n) of three was euthanized per rules set by The University of Georgia Institutional Animal Care and Use Committee. In this study, every ferret that reached humane endpoints exhibited both lethargy (n=1) and dyspnea (n=2) (total score of n=3).

### Hemagglutination inhibition (HAI) assay

The hemagglutination inhibition (HAI) assay was used to quantify HA-specific antibodies by measuring the inhibition in the agglutination of turkey erythrocytes. The protocol was adapted from the WHO laboratory of influenza surveillance manual (33). To inactivate nonspecific inhibitors, the sera was treated with receptor-destroying enzyme (RDE) (Denka Seiken, Co., Japan) prior to being tested. Briefly, three parts RDE was added to one-part sera and incubated overnight at 37°C. RDE was inactivated by incubating at 56°C for approximately 45 minutes. After the incubation period, six parts PBS was added to the RDE-treated sera. RDE-treated sera were two-fold serially diluted in V-bottom microtiter plates. An equal volume of each virus-like particle (VLP) was adjusted to approximately 8 hemagglutination units (HAU)/25 μL, was added to each well. The plates were covered and incubated at RT for 20 minutes before 50 μL RBCs were allowed to settle for 30 minutes at RT.

The HAI titer was determined by the reciprocal dilution of the last well that contained non-agglutinated RBCs. Positive and negative serum controls were included on each plate. Seroprotection was defined as HAI titer of ≥ 1:40 and seroconversion as a 4-fold increase in titer compared to baseline, as defined by the WHO to evaluate influenza vaccines (33).

### Determination of viral nasal wash titers

The nasal washes were performed on anesthetized ferrets by washing out each of their nostrils with a total of 3mL of PBS on days one, three, five and seven post-infection. From each nasal wash, ~2.0mL was recovered. The nasal washes were aliquoted into microcentrifuge tubes and stored at −80°C. Nasal wash aliquots were thawed at RT. Once thawed, 10-fold serial dilutions of nasal washes were overlaid on MDCK cells (33). The MDCK cells were at 95-100% confluency at the time that the assay was performed. Nasal wash samples were incubated for 60 minutes at RT with agitation every 15 minutes. After 60 minutes, the serial dilutions were removed, and the MDCK cells were washed with incomplete (no FBS) DMEM containing Penicillin and Streptomycin (P/S). The wash medium was removed and replaced with 1 mL of a mixture of plaque media without FBS, TPCK-trypsin and 1.2% avicel. Plaque media was made using MEM media, HEPES buffer, L-Glutamine, and P/S. All of the components of the plaque media and avicel were obtained from ThermoFisher (Scientific, Waltham, MA, USA). The MDCK cells were incubated at 37°C with 5% CO_2_ for 48 hours. After 48 hours, the avicel overlay was removed and the MDCK cells were fixed with 10% buffered formalin for a minimum of 15 min. The formalin was then discarded, and the MDCK cells were stained using 1% crystal violet. The MDCK cells were then washed with distilled water to remove the crystal violet. Plaques were then counted and PFU per mL titer was calculated using the number of plaques and the appropriate dilution factor.

### Neutralization Assays

The neutralization assay was used to identify the presence of virus-specific neutralizing antibodies. The protocol was adapted from the WHO laboratory of influenza surveillance manual (33). Equal amounts of sera from each ferret within a vaccination group were combined and heat-inactivated for 30 min at 56 °C. MDCK cells were grown in a 96-well flat bottom plate until they had reached 95-100% confluency. Antibodies were diluted in half-log increments with serum-free media and incubated with 100XTCID_50_ for 1h. The antibody-virus mixture was then added to the incomplete (FBS-free) DMEM washed MDCK cells in the 96-well plate. After 2h, the MDCK cells were washed with incomplete DMEM. Approximately 200 uL of DMEM with P/S and 2.0. ug/mL of TPCK was added to each of the 96 wells. The cell monolayers in the back-titration control wells were checked daily until cytopathic effect (CPE) had reached the majority of the 1XTCID_50_ rows. After three or four days, 50uL of media per well was removed and used in an HA assay to identify the presence of influenza virus. The remaining media in each well was removed and, the MDCK cells were then fixed with 10% buffered formalin for a minimum of 15 min. The formalin was then discarded and the fixed cells were washed with 1x PBS. Afterwards, the MDCK cells were stained using 1% crystal violet (ThermoFisher Scientific, Waltham, MA, USA). The MDCK cells were then washed with distilled water to remove the crystal violet. Any well having an HA activity of ≥1:2 was defined as positive for the analysis. HA activity was confirmed by >10% of CPE in wells that was positive for HA activity.

### Statistical Analysis

Statistical significance was defined as a p-value of less than 0.05. Limit of detection for viral plaque titers was 50 pfu/mL for statistical analysis. The viral plaque titers were transformed by log10 for analysis. The limit of detection for HAI was <1:10 and 1:5 was used for statistical analysis. The HAI titers were transformed by log2 for analysis and graphing. The geometric mean titers were calculated for neutralization assays, but the log2 titers were used for ANOVA analysis. All error bars on the graphs represent standard mean error. ANOVAs with Dunnet’s test was used for weight loss with a statistical significance defined as a p-value of less than 0.05. Nasal wash titers stratified by day and pre-immunity were analyzed with a one-way ANOVA with Tukey Honest Significance Differences method to determine differences between vaccine groups. The overall performance of the vaccines was assessed through multivariate ANOVAs for main effects conducted individually for the neutralization titer, viral nasal wash titer, and HAI titer outcomes followed by Tukey Honest Significance Differences method for adjusting for multiple comparisons. Significantly different groups per outcome was determined from the multiple comparisons. Day 7 of the nasal wash titer was not included in the ANOVA analysis since all of the observations were below the limit of detection. All of the statistical analysis for the various assays can be found in supplemental figures 2–5 and supplemental table S1.

### Amino Acid Sequences

The amino acid sequences for the two COBRA HA sequences have been reported in United States provisional patent filing 14332088_1.

## Results

### Vaccination of ferrets with pre-existing influenza virus immunity

Fitch ferrets (n=20) were made pre-immune with one of three influenza virus subtypes. The H2N2 pre-immunity virus used for infection was either Chk/PA/04 or Qu/RI/16. The H3N2 virus used for infection was either Sich/87 or Pan/99. The H1N1 viruses used for infection were both Sing/86 and CA/09 to represent both seasonal and pandemic H1N1 influenza viruses. After each influenza virus infection, the ferrets were allowed to recover for at least sixty days. Approximately sixty days after the final infection, ferrets had seroconverted to the infection strains with an average HAI titer greater than 1:40 (Fig. S6). The H1N1 alone and H3N2-H1N1 and H1N1-H3N2 pre-immune groups were then infected with their second virus and allowed to recover for an additional sixty days. The H1N1-H3N2 and the H3N2-H1N1 pre-immune groups were then infected with their third virus and allowed to recover for an additional sixty days.

Sixty days after the pre-immune groups’ final viral infection, the ferrets were vaccinated with 15ug of either wild-type (Mal/NL/01 or Mal/WI/08) or COBRA (Z1 or Z5) H2 rHA proteins (Fig. 1). A comparison of the amino acids in the antigenic sites of wild-type and COBRA HA sequences is shown in Fig S1. Four ferrets from each of the pre-immune groups received the same vaccine (five vaccines with four ferrets each equates to twenty ferrets per pre-immunity group). The mock vaccinated group was vaccinated with PBS and adjuvant. Four weeks after the initial vaccination, the ferrets were vaccinated again with 15ug of the same antigen as the first vaccination. All of the ferrets were bled two weeks after each of the vaccinations.

**Figure 1:**
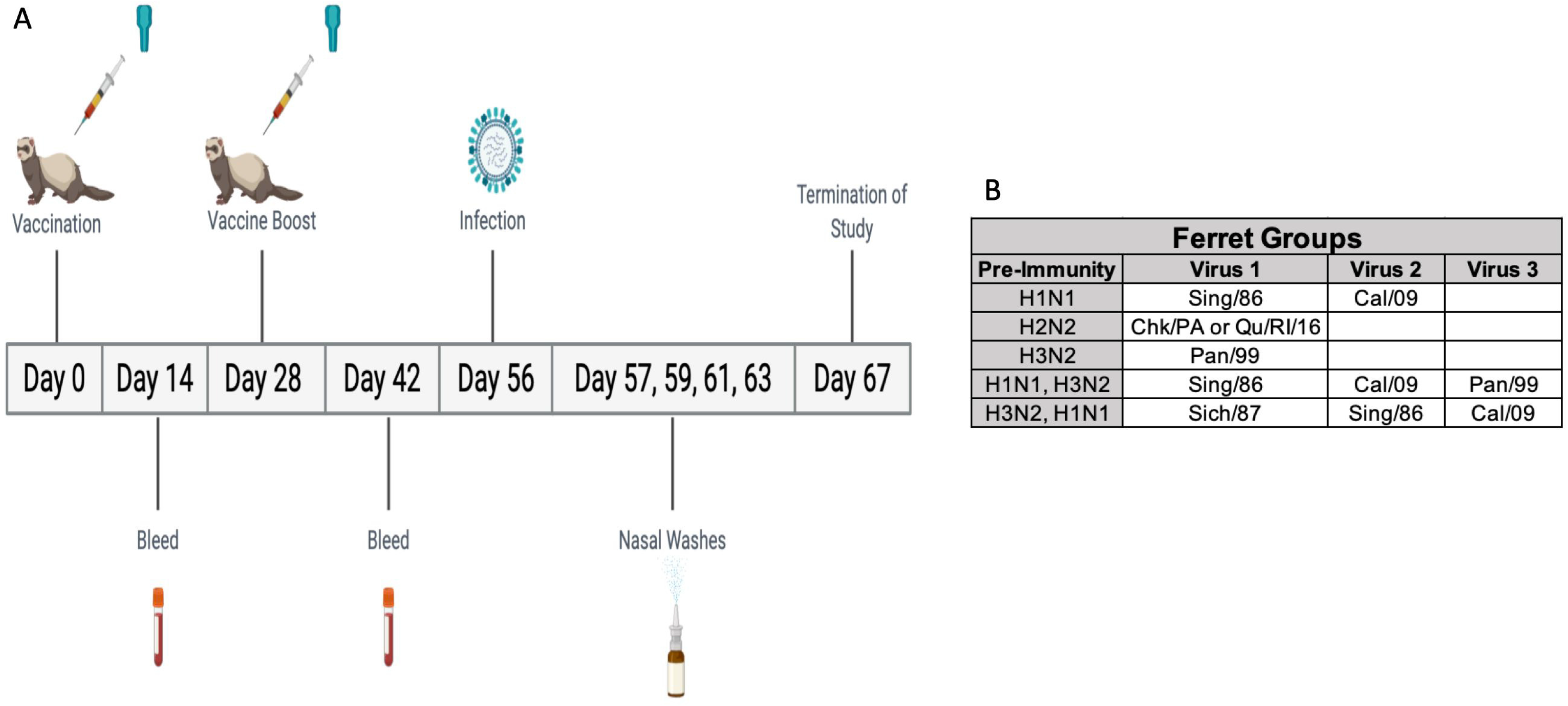
Study Outline. A timeline of the procedures following the establishment of pre-immunity are listed in panel A. Ferrets were vaccinated a D0 and D28. Bleeds were taken at D14 and D42. Ferrets were then infected on D56 and nasal washes were taken on D57, D59, D61, and D63. The study was terminated on D67. The viruses used to establish different pre-immunities are listed in panel B.

### Weight Loss and Survival

Four weeks after the second vaccination, ferrets were infected with the H2N3 clade-3, Sw/MO/06 virus (1e+6 PFU/mL) (Fig. 1). In the H2N2 pre-immune group, there was no significant weight loss after viral challenge and none of the ferrets died (Fig. 2A). In the H3N2 pre-immune group, the ferrets in each of the different H2 rHA vaccination groups had an average weight loss of less than 5% while the mock vaccinated H3N2 pre-immune ferrets reached a peak of 8% average weight loss on D4 post-infection (Fig 2B). The Z5 vaccinated ferrets in the H1N1 pre-immune group reached a peak average weight loss of 6% on D5 post-infection. None of the other vaccination groups (Mal/NL/01 Mal/WI/08, or Z1 COBRA) had an average weight loss of greater than 5% in the H1N1 pre-immune group (Fig. 2C). The H3N2-H1N1 pre-immune ferrets as well as the H1N1-H3N2 pre-immune ferrets had no detectable average weight loss across any of the four vaccination groups or the mock vaccinated group after challenge (Fig. 2D-E).

**Figure 2:**
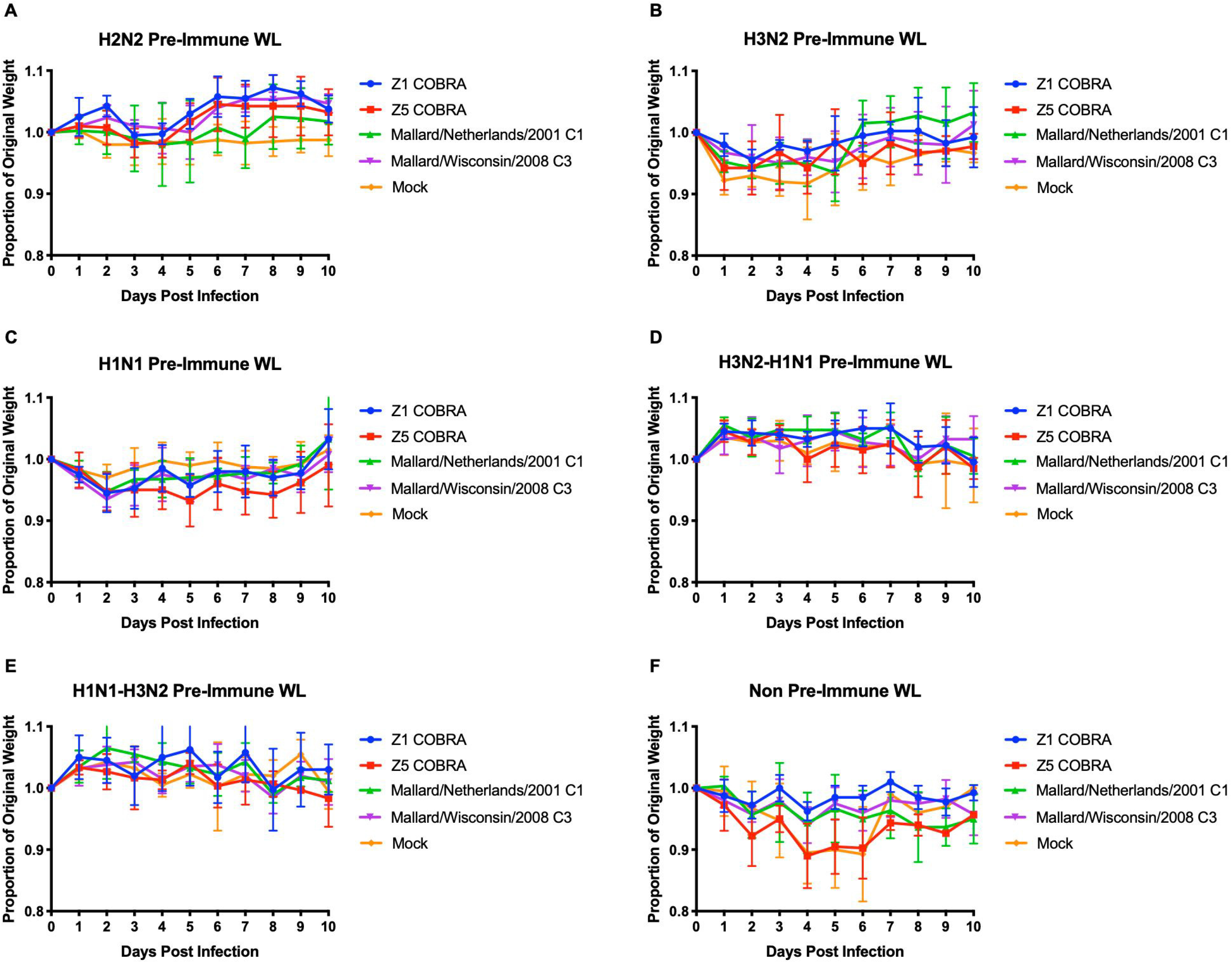
Weight Loss of Viral Challenged Ferrets. Pre-immune ferrets vaccinated with wild-type or COBRA rHA vaccines were challenged with Sw/MO/06. Average weight loss was recorded for each vaccine up to 10 days after infection in the H2N2 pre-immunity (panel A), H3N2 pre-immunity (panel B), H1N1 pre-immunity (panel C), H3N2-H1N1 pre-immunity (panel D), H1N1-H3N2 pre-immunity (panel E), and non pre-immune (panel F) groups. Error bars represent standard mean error.

The non pre-immune ferrets did not have any influenza infection prior to vaccination. Both the Z5 and mock vaccinated ferrets reached a peak average weight loss of >10% by D4 post-infection (Fig. 2F). The Mal/NL/01 and Mal/WI/08 vaccinated groups reached a peak average weight loss of ~5% on D4 post-infection. The Z1 vaccinated group had a peak average weight loss of ~3% on D4 post-infection. The Mal/NL/01, Mal/WI/08, and Z1 vaccinated ferrets all survived until the end of the study. The mock and Z5 vaccinated groups had significantly more weight loss than the Z1 vaccinated ferrets on D4, D5 and D6 (p<0.01 for mock and p<0.05 for Z5). The mock and Z5 vaccinated groups also had significantly more weight loss than the Mal/WI/08 vaccinated ferrets on D5 (p<0.05). These were the only statistically significant differences in weight loss between vaccination groups in any of the pre-immune groups.

Only the non pre-immune and H3N2 pre-immune groups experienced mortality. For the non pre-immune group, one of the Z5 vaccinated ferrets reached humane endpoint by D6 post-infection (Table 1). Also in the non pre-immune group, three of the four ferrets in the mock vaccinated group reached humane endpoint by D6 post-infection (Table 1). In the H3N2 pre-immune group, one ferret in the mock vaccinated group had to be euthanized on D5 post-infection (Table 1).

**Table 1:**
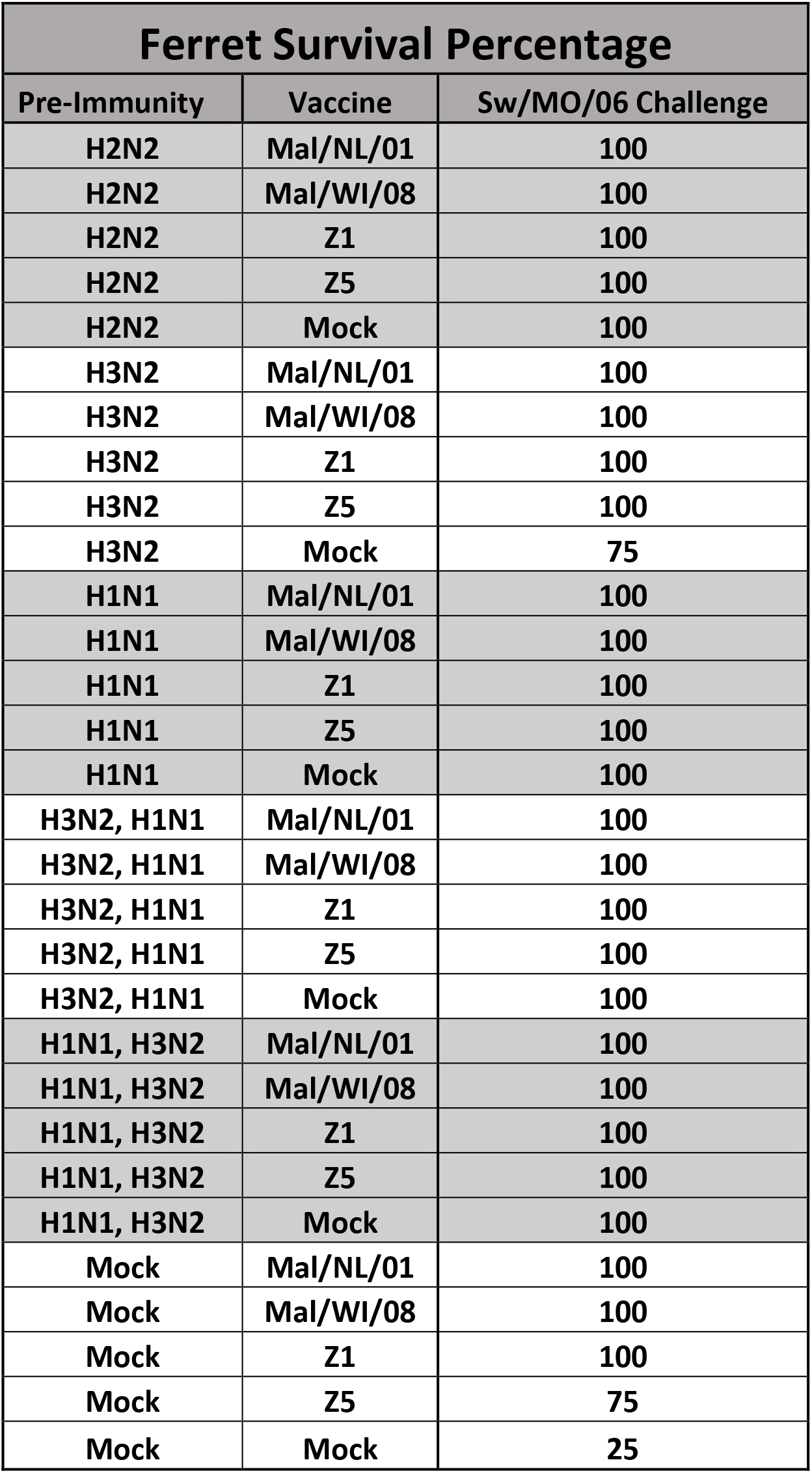
Ferret Survival Percentage. The survival rate among ferrets in each of the vaccination groups across all of the pre-immunities. The columns correspond to the pre-immunity, the vaccine and the survival percentage. Each row is a vaccination group with its pre-immunity followed by the survival percentage following viral challenge. Ferret groups either contained n=3 or n=4 ferrets.

### Viral Titers

In the H2N2 pre-immune group, one ferret in both the Z5 and mock vaccinated groups had viral titers of ~1.0e+3 PFU/mL on D1. One ferret each of the Mal/NL/01, Z5, and mock vaccinated groups had detectable viral titers in their nasal washes at D3. There were no detectable viral titers in any ferret in the D5 or D7 nasal washes (Fig. 3A-D). In the H3N2 pre-immune group, one ferret in each of the Mal/NL/01, Z1 and Z5 vaccination groups had viral titers on D1 (between 1e+2 and 1e+3) (Fig. 3E). The mock vaccinated group had two ferrets with viral titers around 1e+4 while the Mal/WI/08 ferrets had no detectable titers. On D3 post-infection, one ferret in each of the vaccination groups besides the mock vaccination group had viral titers between 5e+2 and 5e+3 PFU/mL. The mock vaccinated ferrets had no detectable titers on D3 (Fig. 3F). On D5 post-infection, only one ferret in the mock vaccinated group had detectable viral titers (Fig 3G). This mock vaccinated ferret also reached humane endpoints on D5 (Table 1). None of the H3N2 pre-immune ferrets had detectable viral titers on D7 post-infection (Fig 3H).

**Figure 3:**
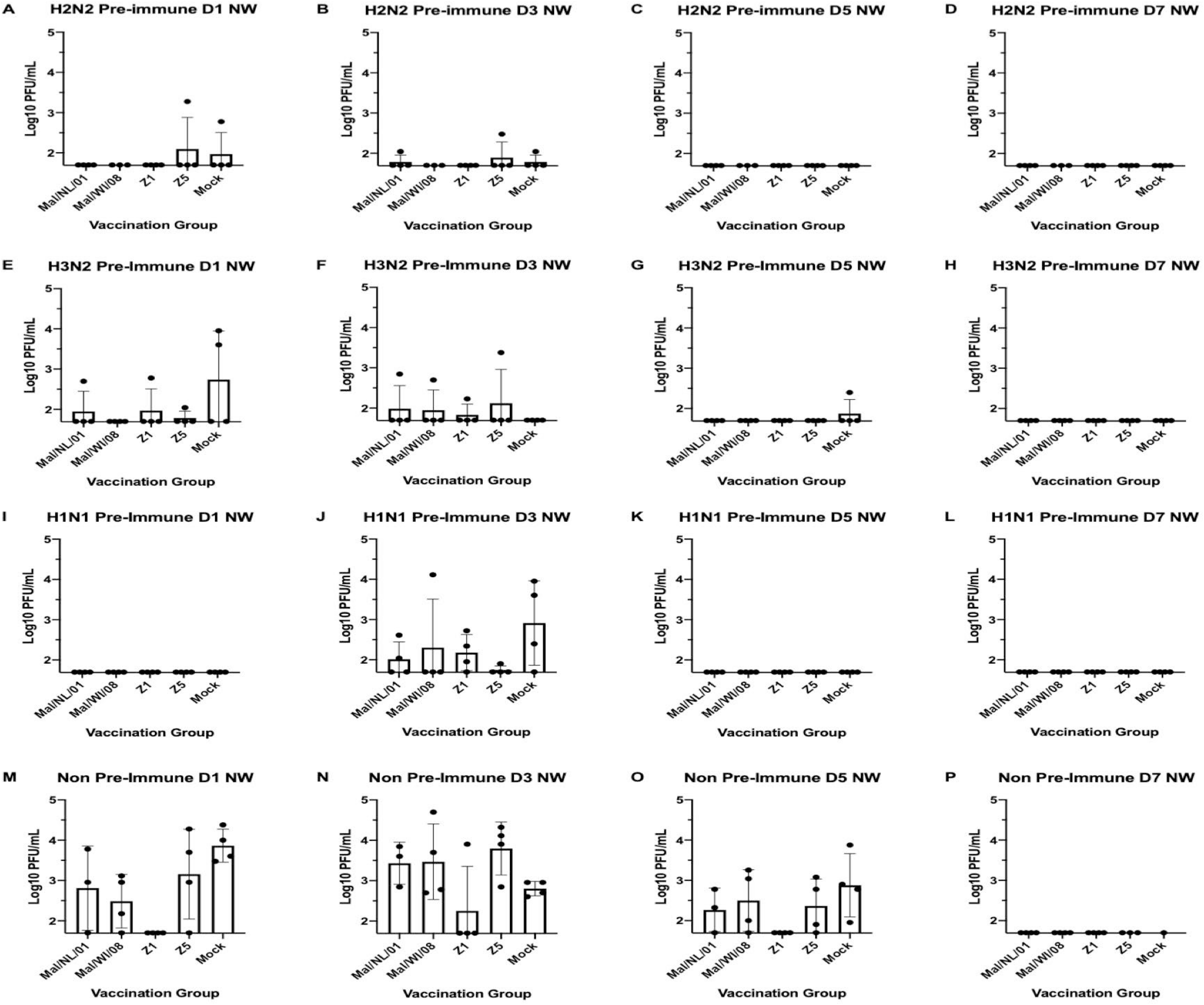
Viral Nasal Wash Titers. Nasal washes were performed on D1, D3, D5 and D7 post-infection. The titers are recorded as log10 PFU/mL. The H2N2 pre-immune ferrets are shown in panels A-D. The H3N2 pre-immune ferrets are shown in panels E-H. The H1N1 pre-immune ferrets are shown in panels I-L. The non pre-immune ferrets are shown in panels M-P. The height of the bars shows the mean while the error bars represent mean standard error.

None of the ferrets in the H1N1 pre-immune group had detectable viral titers in their nasal washes on D1, D5, or D7 post-infection (Fig. 3I, 3K-L). All five of the vaccination groups had at least one ferret with detectable viral titers in their nasal washes on D3. The mock vaccinated ferrets had the highest average viral titers (~1e+3 PFU/mL) while the other four vaccination groups had average viral titers between 5e+1 and 5e+2 PFU/mL (Fig. 3J).

In the non pre-immune group, multiple ferrets in Mal/NL/01, Mal/WI/08, Z5, and mock vaccination groups all had detectable viral titers in their D1 nasal washes with multiple ferrets in each of the vaccination groups having ≥3e+1 viral titers. None of the ferrets in the Z1 vaccination group had detectable viral titers on D1 post-infection (Fig. 3M). On D3 post-infection, the average viral titers for the Mal/NL/01, Mal/WI/08, Z5, and mock vaccinated ferrets all had average viral titers between ~1e+3 and 1e+4. One ferret in the Z1 vaccination group had detectable viral titers on D3 post-infection (Fig. 3N). On D5 post-infection, multiple ferrets in the Mal/NL/01, Mal/WI/08, Z5, and mock vaccinated groups had viral titers >1e+2 with the mock vaccinated ferrets having the highest average viral titer of ~1e+3. None of the ferrets in the Z1 vaccination group had detectable viral titers on D5 post-infection (Fig. 3O). None of the surviving ferrets had detectable viral titers in their nasal washes on D7 post-infection (Fig. 3P). None of the ferrets in the H3N2-H1N1 pre-immune or the H1N1-H3N2 pre-immune groups had any detectable viral titers in their nasal washes on any day (Fig. S7). Of all the time points and pre-immunity background there was no significant difference between the vaccinated groups and the mock groups, except for non pre-immune D1 NW, where Z1 was significantly lower than the mock group with p-value of 0.0078, with a one-way ANOVA + Tukey.

A three-way ANOVA looking at the main effects of vaccine received, the ferret pre-immunity, and the day of the nasal wash, indicated that overall when adjusting for pre-immunity and day post infection that the mean viral nasal wash titers of the Z1 COBRA was significantly lower than that of the mock vaccinated group by 0.322 log 10 viral titer (p adjusted < 0.001). Furthermore, the Z1 COBRA also had a titer 0.219 log 10 lower after adjustment compared to the Z5 COBRA (p adjusted = 0.039) (Fig S2). Only the non pre-immune ferrets were significantly different from the other pre-immunities after controlling for vaccine received and day post infection (Fig S2). All other pre-immunities have nonsignificant mean viral titers. When comparing the day post infection, Day 1 and day 3 were not significantly different, but day 5 had lower viral titers compared to either day 3 or day 1.

### Hemagglutination-inhibition (HAI) antibodies

The HAI titers varied greatly between the pre-immune groups. The H2N2 pre-immune ferrets were the only pre-immune group to have HAI titers to VLPs in the H2 panel on the day of prime vaccination (Fig. 4). Ferrets in all five of the vaccination groups had average HAI titers above 1:40 to multiple VLPs in the H2 panel, but none of the vaccination groups had detectable titers to all 12 of the H2 HA VLPs (Fig. 4).

**Figure 4:**
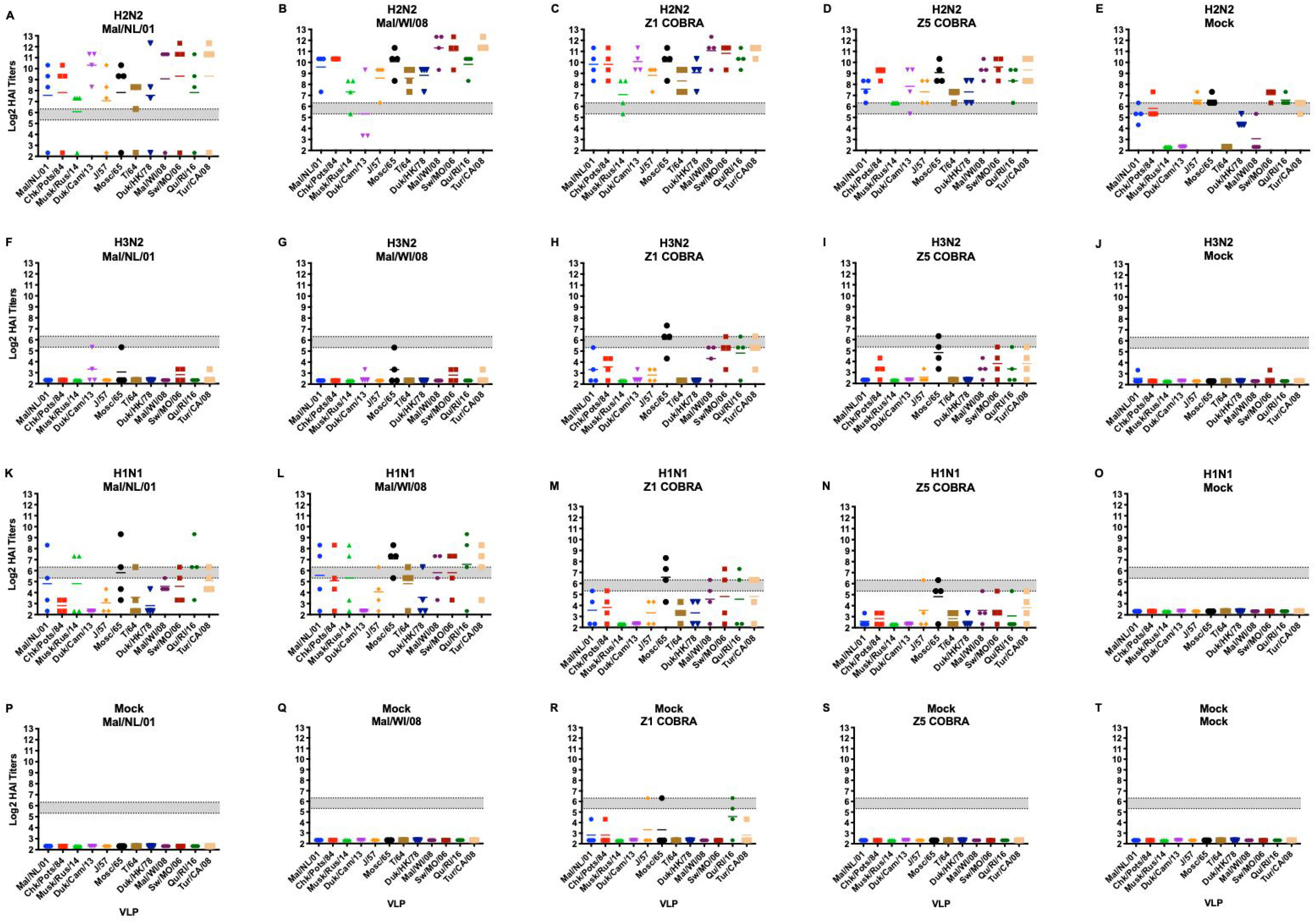
Antibody Cross-Reactive Antibody Responses on D0 (Prime Vaccination) in H2N2 Pre-immune Ferrets. HAI titers for each vaccine group in the H2N2 pre-immunity. Serum from each ferret was obtained on the day of prime vaccination (sixty days post-infection) and tested against VLPs expressing 12 WT H2 HA sequences. Dotted lines indicate 1:40 and 1:80 HAI titer respectively. The VLP panel is composed of: clade-1 HAs (Mallard/Netherlands/2001, Chicken/Potsdam/1984, Muskrat/Russia/2014, Duck/Cambodia/2013), clade-2 HAs (Duck/Hong Kong/1978, Taiwan/1/1964, Moscow/1019/1965, Japan/305/1957), and clade-3 HAs (Mallard/Wisconsin/2008, Swine/Missouri/2006, Quail/Rhode Island/2016, Turkey/California/2008). Error bars represent standard mean error.

After the first vaccination, the H2N2 pre-immune ferrets had a geometric mean HAI titer of ≥1:40 to all twelve of the VLPs in the panel excluding the mock vaccination group (Fig. 5A-E). The H3N2 pre-immune ferrets had low HAI titers to the H2 VLPs after the first vaccination (Fig. 5F-J). Only the Z1 vaccinated ferrets had a geometric mean HAI titer of ≥1:40 to more than one H2 VLP (Fig. 5H). The mock vaccinated ferrets in H1N1 pre-immune group did not have detectable HAI titers to any of the H2 VLPs after the first vaccination. The Mal/NL/01, Mal/WI/08, and Z1 vaccinated ferrets had geometric mean HAI titers of ≥1:40 against multiple H2 VLPs after the first vaccination (Fig. 5K-L, N-O). The Z5 vaccinated H1N1 pre-immune ferrets only had geometric mean HAI titers of ≥1:40 against one H2 VLP (Fig. 5M). The non-pre-immune ferrets in the Mal/NL/01, Mal/WI/08, Z5, and mock vaccination groups had no detectable HAI titers after the first vaccination (Fig. 5P-Q, S-T). The Z1 vaccinated ferrets had some HAI titers after the first vaccination, but none of them reached a geometric mean titer of 1:40 (Fig. 5R).

**Figure 5:**
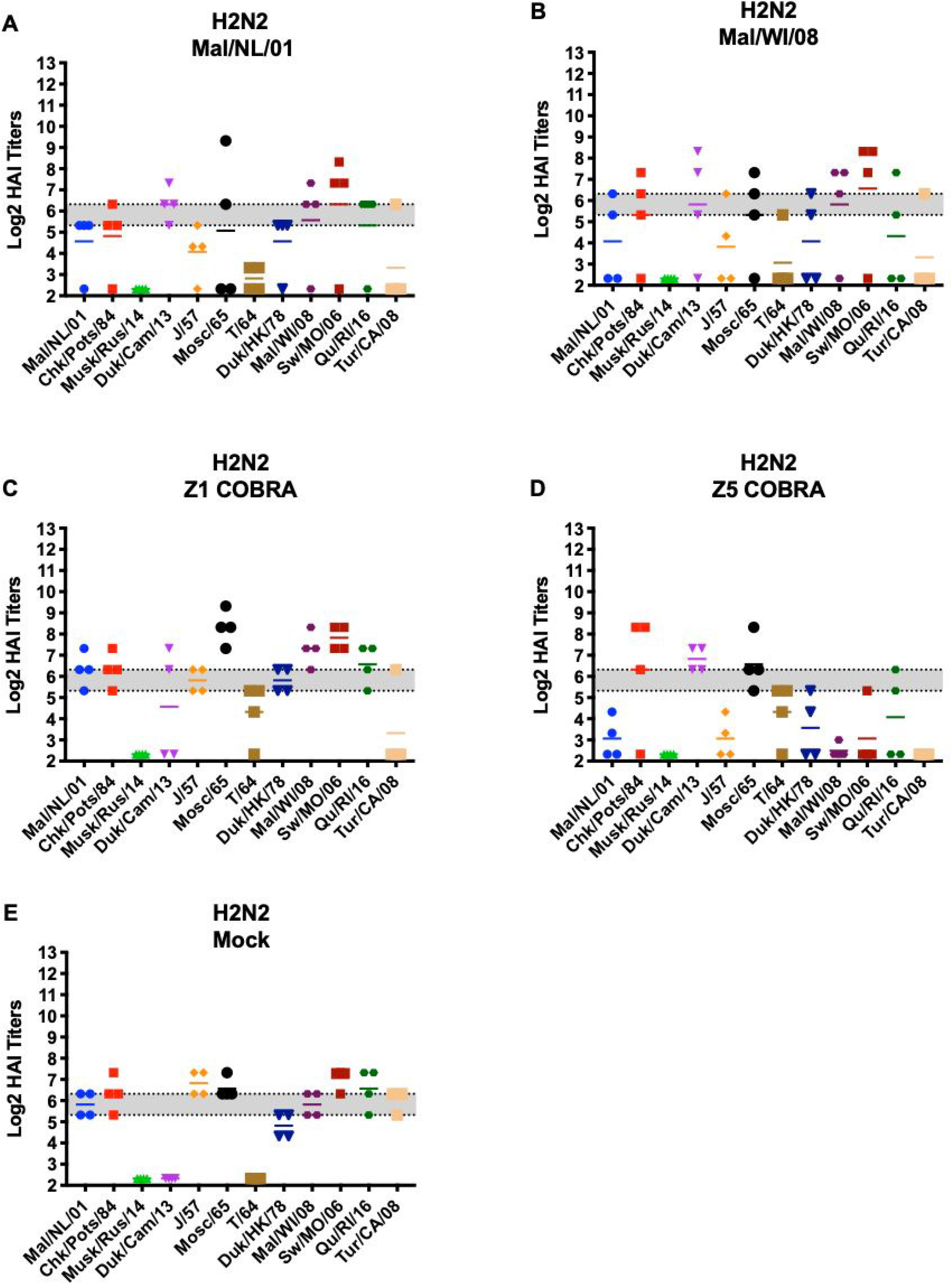
Antibody Cross-Reactivity of H2N2, H1N1, H3N2 and non pre-immune Groups on D14 post-prime vaccination. HAI titers for each vaccine group 14 days after the first vaccine in the H2N2 pre-immunity (panels A-E), H3N2 pre-immunity (panels F-J), H1N1 pre-immunity (panels K-O), and non pre-immune (panels P-T) groups. Serum from each ferret was obtained on day 14 after the first vaccination and tested against VLPs expressing 12 WT H2 HA sequences. Dotted lines indicate 1:40 and 1:80 HAI titer respectively. The VLP panel is composed of: clade-1 HAs (Mallard/Netherlands/2001, Chicken/Potsdam/1984, Muskrat/Russia/2014, Duck/Cambodia/2013), clade-2 HAs (Duck/Hong Kong/1978, Taiwan/1/1964, Moscow/1019/1965, Japan/305/1957), and clade-3 HAs (Mallard/Wisconsin/2008, Swine/Missouri/2006, Quail/Rhode Island/2016, Turkey/California/2008). Error bars represent standard mean error.

After the second vaccination, the HAI titers for the ferrets in the H2N2 pre-immune group did not drastically change (Fig. 6A-E). In the H3N2 pre-immune group, HAI titers in the Mal/NL/01, Mal/WI/08, Z1 and Z5 vaccinated ferrets all increased significantly after the second vaccination. This was the only statistically significant increase in average titer using multiple t-test analysis comparing for the change in each vaccine titer between days 14 and 42 post-prime vaccination. Each of these four vaccination groups had HAI titers of ≥1:40 to seven or more VLPs in the panel. The Z1 and Z5 vaccinated ferrets had geometric mean HAI titers of ≥1:80 to nine and ten of the twelve VLPs in the panel respectively (Fig. 6F-J).

**Figure 6:**
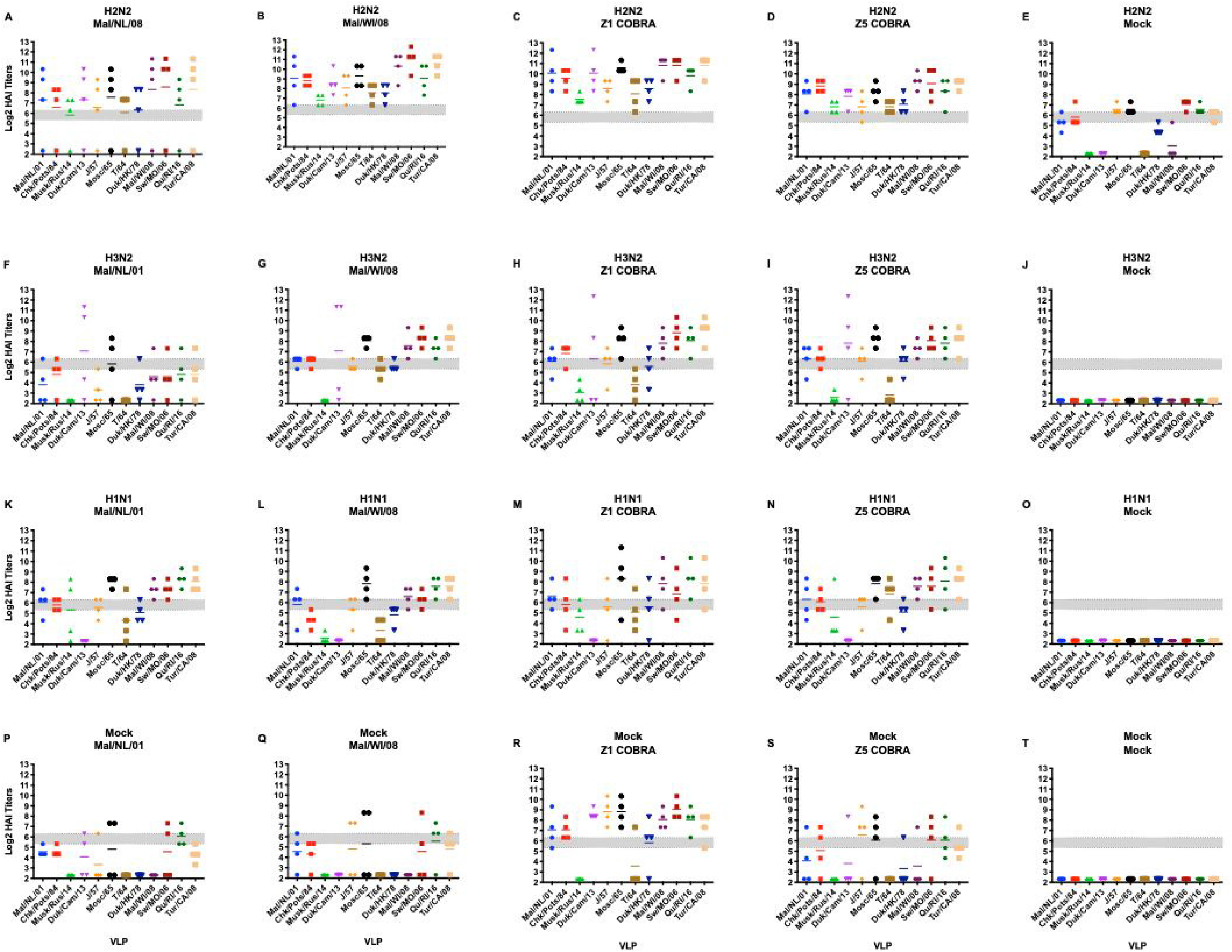
Antibody Cross-Reactivity of H2N2, H1N1, H3N2 and non pre-immune Groups D42 post-prime (D14 post-boost) vaccination. HAI titers for each vaccine group 14 days after the second vaccination in the H2N2 pre-immunity (panels A-E), H3N2 pre-immunity (panels F-J), H1N1 pre-immunity (panels K-O), and non pre-immune (panels P-T) groups. Serum from each ferret was obtained on day 42 post-vaccination and tested against VLPs expressing 12 WT H2 HA sequences. Dotted lines indicate 1:40 and 1:80 HAI titer respectively. The VLP panel is composed of: clade-1 HAs (Mallard/Netherlands/2001, Chicken/Potsdam/1984, Muskrat/Russia/2014, Duck/Cambodia/2013), clade-2 HAs (Duck/Hong Kong/1978, Taiwan/1/1964, Moscow/1019/1965, Japan/305/1957), and clade-3 HAs (Mallard/Wisconsin/2008, Swine/Missouri/2006, Quail/Rhode Island/2016, Turkey/California/2008). Error bars represent standard mean error.

In the H1N1 pre-immune ferrets, the HAI titers of all of the vaccination groups besides the mock vaccination group increased after the second vaccination (Fig. 6K-O). After the second vaccination, the Mal/WI/08 vaccination group had geometric mean HAI titers of ≥1:40 to seven of the twelve VLPs in the panel. The Mal/NL/01, Z1 and Z5 vaccinated ferrets all had geometric mean HAI titers of >1:40 to ten or more of the VLPs in the H2 panel. The Mal/NL/01 and the Mal/WI/08 vaccinated ferrets had a geometric mean HAI titer of ≥1:80 to seven and six of the twelve VLPs in the H2 panel respectively. The Z1 and Z5 vaccinated ferrets had geometric mean HAI titers of ≥1:80 to nine of the twelve VLPs in the H2 panel.

In the non pre-immune group, the HAI titers in the Mal/NL/01, Mal/WI/08, Z1 and Z5 vaccination groups all increased after the second vaccination (Fig. 6P-T). The Mal/NL/01 and Mal/WI/08 vaccinated ferrets had geometric mean HAI titers of ≥1:40 to three and five of the VLPs in the H2 panel. The Z1 and Z5 vaccinated ferrets had geometric mean HAI titer of ≥1:40 to eleven and nine of the VLPs respectively. The Mal/NL/01, Mal/WI/08, and Z5 vaccinated ferrets had geometric mean HAI titer of ≥1:80 to two, four, and five of the H2 VLPs in the panel. The Z1 vaccinated ferrets had geometric mean HAI titers of ≥1:80 to ten of the twelve VLPs in the H2 panel.

The H3N2-H1N1 and the H1N1-H3N2 pre-immune groups, varied greatly in their HAI responses. The H3N2-H1N1 pre-immune group had little to no HAI titers after the first vaccination to any of the VLPs in the panel (Fig. 7A-E). The H1N1-H3N2 pre-immune group had multiple ferrets in each vaccination group with detectable HAI titers to each of the twelve VLPs in the panel (Fig 7F-J). The Z5 and Mal/NL/01 vaccination groups had a geometric mean HAI titer of >1:40 to seven and eight of the VLPs in the panel, respectively. The Mal/WI/08 and Z1 vaccination groups had a geometric mean HAI titer of ≥1:40 to eleven and twelve out of the twelve VLPs in the panel, respectively.

**Figure 7:**
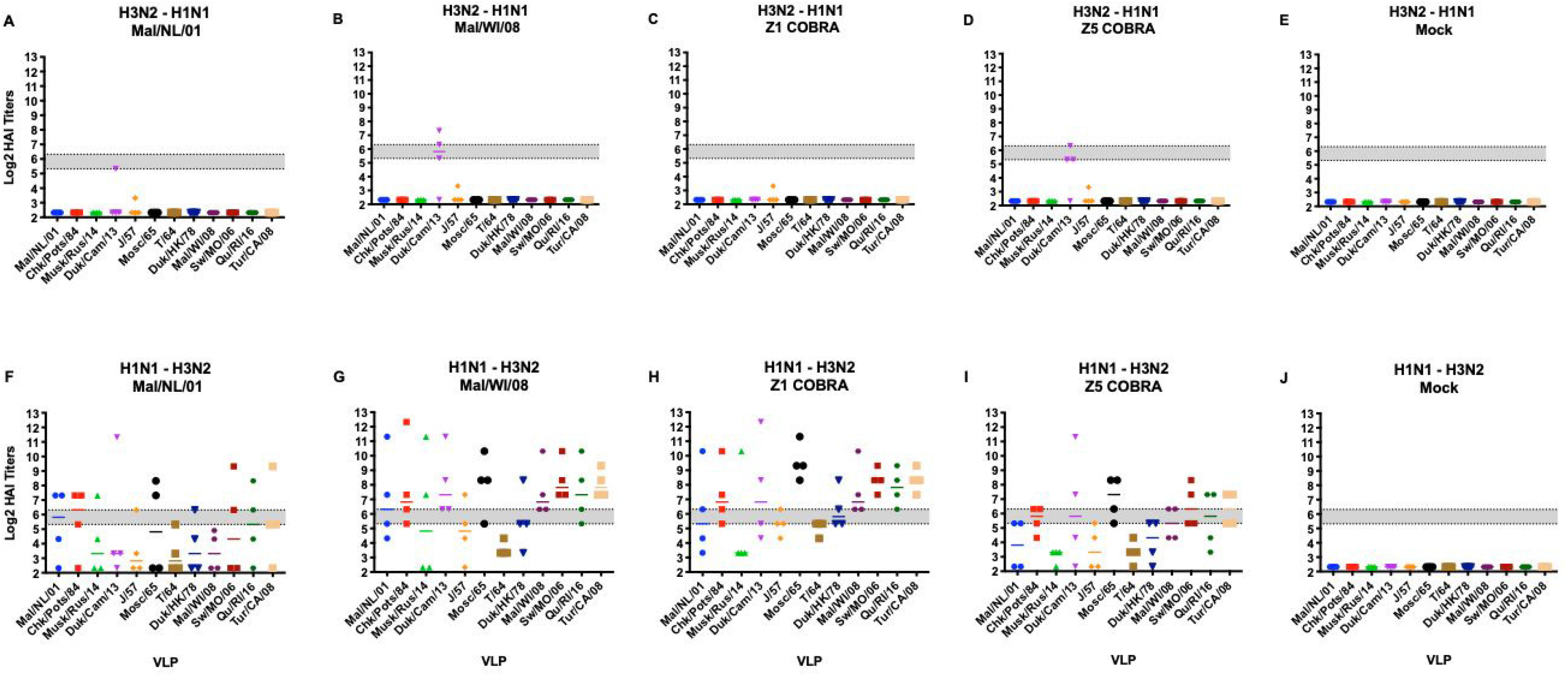
Antibody Cross-Reactivity of H1N1-H3N2, and H3N2-H1N1 pre-immune groups D14 post-prime vaccination. HAI titers for each vaccine group 14 days after the first vaccination (D14) in the H1N1-H3N2 pre-immunity (panels A-E), H3N2-H1N1 pre-immunity (panels F-J) groups. Dotted lines indicate 1:40 and 1:80 HAI titer respectively. The VLP panel is composed of: clade-1 HAs (Mallard/Netherlands/2001, Chicken/Potsdam/1984, Muskrat/Russia/2014, Duck/Cambodia/2013), clade-2 HAs (Duck/Hong Kong/1978, Taiwan/1/1964, Moscow/1019/1965, Japan/305/1957), and clade-3 HAs (Mallard/Wisconsin/2008, Swine/Missouri/2006, Quail/Rhode Island/2016, Turkey/California/2008). Error bars represent standard mean error.

After the second vaccination, the Z5 and Z1 vaccinated ferrets in H3N2-H1N1 pre-immune group had HAI titers of ≥1:40 to five and six of the VLPs in panel, respectively. The Mal/NL/01 and the Mal/WI/08 vaccinated ferrets had HAI titers of ≥1:40 to nine and ten of the twelve VLPs in the panel, respectively (Fig. 8A-E). After the second vaccination for the H1N1-H3N2 pre-immune group, the Z5 vaccinated ferrets had a geometric mean HAI titer of ≥1:40 to ten of the VLPs while the Z1 vaccinated ferrets had a geometric mean HAI ≥1:40 to all twelve of the VLPs in the panel. The Mal/NL/01 and Mal/WI/08 vaccination groups had a geometric mean HAI titer of ≥1:40 to eleven of the VLPs in the panel (Fig. 8F-J).

**Figure 8:**
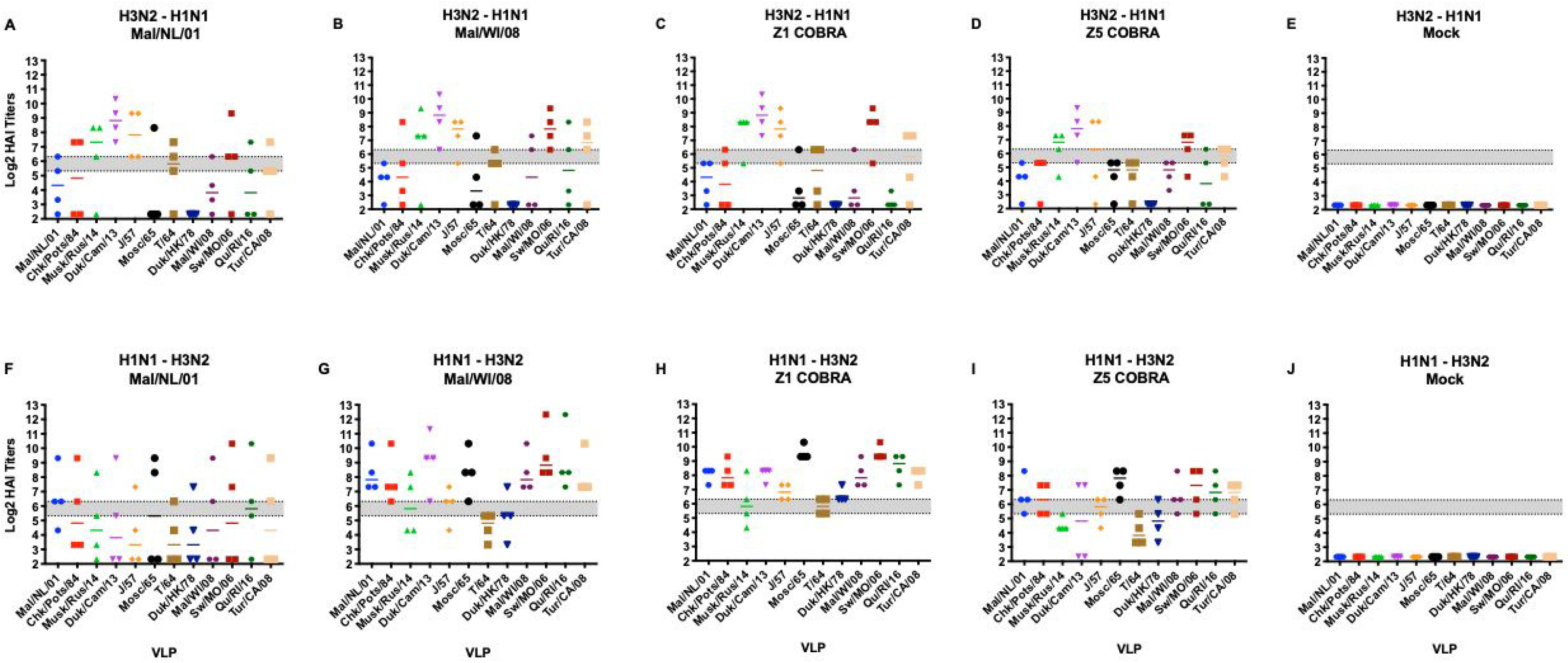
Antibody Cross-Reactivity of H1N1-H3N2, and H3N2-H1N1 pre-immune groups D42 post-prime (D14 post-boost) vaccination. HAI titers for each vaccine group 14 days after the second vaccination (D42) in the H1N1-H3N2 pre-immunity (panels A-E), H3N2-H1N1 pre-immunity (panels F-J) groups. Dotted lines indicate 1:40 and 1:80 HAI titer respectively. The VLP panel is composed of: clade-1 HAs (Mallard/Netherlands/2001, Chicken/Potsdam/1984, Muskrat/Russia/2014, Duck/Cambodia/2013), clade-2 HAs (Duck/Hong Kong/1978, Taiwan/1/1964, Moscow/1019/1965, Japan/305/1957), and clade-3 HAs (Mallard/Wisconsin/2008, Swine/Missouri/2006, Quail/Rhode Island/2016, Turkey/California/2008). Error bars represent standard mean error.

After controlling for the main effects of vaccine received, preexisting immunity and HAI VLP, the Z1 COBRA vaccine had a significantly higher overall log2 mean HAI titer compared to the other vaccine groups using an ANOVA with Tukey adjustment method. The Z5 COBRA and Mal/WI/08 vaccines were not significantly different from each other, and the Mal/NL/01 vaccine titers were significantly lower than the other vaccine groups. For pre-immunity, the ferrets with an H2N2 background had a significantly higher mean titer than compared to the H1N1, H3N2, and H1N1-H3N2 groups which were not significant from each other. The H3N2-H1N1 pre-immunity overall mean was not significantly different from the mock pre-immunity mean titer.

### Neutralization Assays

Serum was collected from the ferrets after the second vaccination and pooled in equal amounts for the neutralization assay. The Mal/NL/01, Mal/WI/08, and Z1 vaccinated ferrets with the H2N2 pre-immunity all had neutralizing antibodies titers ≥1:350 to five of the seven influenza viruses in the neutralization panel (Table 2). The Z5 vaccinated ferrets had neutralizing antibodies titers ≥1:350 to six of the seven influenza viruses. The H3N2 pre-immune ferret groups all had neutralization titers lower than 1:200 to all of the influenza viruses in panel except for Sw/MO/06 where the Mal/NL/01, Mal/WI/08, Z1, and Z5 vaccinated ferrets all had titers >1:450. In the H1N1 pre-immune group, the Mal/NL/01 and Z5 vaccine groups had neutralization titers >1:100 to two of the seven influenza viruses in the panel while the Mal/WI/08 and Z1 vaccine groups had neutralization titers >1:100 for four and five of the influenza viruses, respectively. The Mal/NL/01 and Z5 vaccinated ferrets in the H3N2-H1N1 pre-immune group had neutralization titers >1:200 to two of the seven influenza viruses in the H2Nx virus panel. The Mal/WI/08 vaccinated ferrets had neutralization titers >1:200 only to the Sw/MO/06 H2N3 influenza virus. The Z1 vaccinated ferrets had titers >200 to five of the seven influenza viruses. For the H1N1-H3N2 pre-immune group, the Mal/NL/01, Mal/WI/08, Z1, and Z5 vaccinated ferrets all had neutralization ferrets >1:150 to four of the seven influenza viruses. For the non-pre-immune group, the Mal/NL/01, Mal/WI/08, Z1, and Z5 vaccinated ferrets all had neutralization titers >1:350 to the Sw/MO/06 influenza virus. The Z1 vaccinated ferrets had neutralization titers >1:150 to three of the six other influenza viruses in the panel while the Z5 vaccinated ferrets had neutralization titers >1:150 to one other influenza virus in the panel.

**Table 2:** Ferret Neutralization Titers on D42 post-prime sera (D14 post-boost) Neutralization titers were obtained from pooled sera. Titers were obtained by taking the geometric mean titer of the replicates for each of the vaccine groups. Each column is a virus that was used in the neutralization assay. Each row is the sera from a vaccination group. The lower limit of detection is 5 while the upper limit of detection is 640. Significance group is determined for each individual pre-immunity group. Comparing within one pre-immune group, the vaccines’ ability to elicit neutralization titers were compared, while controlling for the main effects of virus used in the assay.

| Ferret Neutralization Titers D42 |  |  |  |  |  |  |  |  |  |
| --- | --- | --- | --- | --- | --- | --- | --- | --- | --- |
| Pre-Immunity | Vaccine | Significance group | Chk/Pots/84 | Chk/PA/04 | For/57 | T/64 | Duk/HK/78 | Sw/MO/06 | Mall/MN/08 |
| H2N2 | Mal/NL/01 | a | 640 | 380 | 48 | 34 | 640 | 640 | 452 |
| H2N2 | Mal/WI/08 | a | 640 | 537 | 48 | 56 | 640 | 640 | 640 |
| H2N2 | Z1 | a | 640 | 640 | 134 | 67 | 640 | 640 | 640 |
| H2N2 | Z5 | a | 452 | 640 | 40 | 380 | 537 | 640 | 320 |
| H2N2 | Mock | b | 95 | 156 | 8 | 24 | 48 | 537 | 95 |
| H3N2 | Mal/NL/01 | b | 17 | 7 | 5 | 20 | 12 | 452 | 17 |
| H3N2 | Mal/WI/08 | ab | 48 | 67 | 5 | 28 | 24 | 640 | 67 |
| H3N2 | Z1 | a | 190 | 56 | 7 | 40 | 67 | 640 | 95 |
| H3N2 | Z5 | ab | 113 | 113 | 5 | 20 | 34 | 640 | 48 |
| H3N2 | Mock | c | 5 | 7 | 5 | 5 | 5 | 6 | 6 |
| H1N1 | Mal/NL/01 | b | 113 | 20 | 5 | 95 | 28 | 537 | 24 |
| H1N1 | Mal/WI/08 | ab | 113 | 40 | 6 | 537 | 34 | 640 | 226 |
| H1N1 | Z1 | a | 537 | 269 | 20 | 80 | 269 | 640 | 269 |
| H1N1 | Z5 | ab | 134 | 48 | 10 | 20 | 80 | 640 | 12 |
| H1N1 | Mock | c | 8 | 6 | 5 | 5 | 5 | 5 | 5 |
| H3N2, H1N1 | Mal/NL/01 | ac | 269 | 160 | 28 | 34 | 95 | 640 | 134 |
| H3N2, H1N1 | Mal/WI/08 | a | 113 | 113 | 10 | 34 | 67 | 640 | 134 |
| H3N2, H1N1 | Z1 | c | 640 | 380 | 28 | 160 | 537 | 640 | 226 |
| H3N2, H1N1 | Z5 | ac | 226 | 160 | 17 | 48 | 160 | 640 | 160 |
| H3N2, H1N1 | Mock | b | 5 | 5 | 7 | 5 | 5 | 6 | 6 |
| H1N1, H3N2 | Mal/NL/01 | a | 640 | 640 | 34 | 5 | 80 | 640 | 640 |
| H1N1, H3N2 | Mal/WI/08 | a | 537 | 269 | 40 | 40 | 80 | 640 | 380 |
| H1N1, H3N2 | Z1 | a | 226 | 640 | 67 | 113 | 95 | 640 | 537 |
| H1N1, H3N2 | Z5 | a | 269 | 269 | 20 | 40 | 113 | 640 | 190 |
| H1N1, H3N2 | Mock | b | 5 | 6 | 5 | 5 | 5 | 5 | 5 |
| Mock | Mal/NL/01 | b | 34 | 14 | 5 | 10 | 10 | 640 | 20 |
| Mock | Mal/WI/08 | ab | 12 | 17 | 5 | 5 | 7 | 640 | 20 |
| Mock | Z1 | c | 537 | 190 | 40 | 80 | 226 | 640 | 80 |
| Mock | Z5 | b | 160 | 28 | 6 | 10 | 56 | 380 | 14 |
| Mock | Mock | a | 5 | 5 | 5 | 5 | 5 | 5 | 5 |

## Discussion

Currently, the United States stockpiles pre-pandemic influenza virus vaccines for both H5 and H7 influenza subtypes (34). While both H5 and H7 influenza viruses have infected humans, there has been little human-to-human transmission of these influenza virus subtypes. Conversely, H2N2 influenza viruses caused the 1957 influenza pandemic (3). It is likely that H2 influenza viruses may cause future pandemics and therefore next-generation, broadly reactive universal influenza vaccines should elicit protective immune responses against current and future H2 influenza viruses. Currently, humans develop long-lasting memory cells against seasonal influenza A and B viruses following infection and/or vaccination. However, few studies have investigated the effects of vaccinating individuals for a novel subtype of influenza virus in the presence of pre-existing immunity to historical influenza strains. This study addressed the effect of vaccinating ferrets with H2 influenza rHA vaccines that have pre-existing immunity induced by historical influenza A virus (IAV) strains and the effectiveness of broadly reactive COBRA H2 rHA vaccines to stimulate protective antibody titers.

IAV naïve ferrets were infected individually with H1N1, H2N2, or H3N2 influenza viruses prior to H2 rHA vaccination. Some ferrets were also infected sequentially with H1N1 and H3N2 influenza viruses. Following a single vaccination of the COBRA HA vaccines, ferrets pre-immune to group 1 influenza A viruses all had broadly cross-reactive antibodies. This included the ferrets that were administered the H1N1 virus prior to the H3N2 virus. Ferrets infected with H3N2 influenza viruses initially had no cross-reactive HAI titers to H2 influenza viruses after the first vaccination, which was similar to naïve ferrets. This result shows that the first subtype of influenza virus that a person is infected with affects their future immune responses to subsequent heterotypic influenza virus vaccinations. This phenomenon, termed ‘Immune Imprinting’, has been well-documented for IAV (26–28). These previous studies showed that the IAV that an individual was first infected with seemed to confer protection to other influenza virus subtypes within the same HA evolutionary group. Group 1 influenza virus HA subtypes include H1, H2, H5 and H9 while group 2 influenza virus HA subtypes include H3 and H7 (27, 28). In this study, the broader HAI activity after the prime vaccination in the H1N1 pre-immune ferrets compared to the H3N2 pre-immune ferrets further support the IAV group specific immune imprinting theory.

*De novo* immune responses appear more important and are required for the H3N2 (Group 2) imprinted ferrets to generate cross-reactive antibodies to H2 influenza viruses than the H1N1 or H2N2 (Group 1) pre-immune ferrets. Once the animals were administered a second vaccination, the H3N2 and H3N2-H1N1 pre-immune groups had detectable HAI titers, but the titers were 2-4 fold lower on average than Group 1 imprinted ferrets. The Group 2 imprinted ferrets likely have B-cells that are highly specific to epitopes on the H3 HA (35). These epitopes are far less similar to the epitopes on H2 HA than the epitopes on H1 HA. Without any similar immune memory, the H3N2 influenza virus imprinted ferrets would likely need to generate *de novo* B cells to the COBRA H2 HA vaccine (27). Meanwhile, the H1N1 influenza virus imprinted ferrets have memory B cells that may recognize similar epitopes on the H2 HA proteins. These memory B cells would undergo somatic hypermutation and adapt to the H2 HA epitopes (36). Therefore, sequential infections of influenza viruses may affect the immune responses to future influenza virus infections or vaccinations (27, 28, 35, 36) Since most people under 55 years old will have pre-existing memory cells to both H1N1 and H3N2 influenza viruses, a stockpiled H2 influenza virus vaccine will likely need a prime and boost vaccination in order to generate *de novo* immune responses to adequately protect the majority of individuals from a future H2 influenza virus infections.

Mortality in the mock vaccinated ferrets was absent in all of the pre-immune groups with the exception of the naïve pre-immune ferrets following the H2N3 virus challenge. This result was surprising since none of the mock vaccinated ferrets in any of the pre-immune groups (with the exception of the H2N2 pre-immune ferrets) had substantial HAI or neutralization titers to any of the H2 VLPs or influenza viruses tested in these assays. Given these results, it is likely that other immune mechanisms may be playing a role in protecting ferrets from mortality during the viral challenge. Without H2 specific neutralizing antibodies, it is possible that either non-neutralizing antibodies or T-cells are contributing to protection against the Sw/MO/06 H2N3 virus infection (37–40). Non-neutralizing antibodies could be contributing to protection through antibody-dependent cellular cytotoxicity (ADCC) or complement-dependent cytotoxicity (41–43). Both CD4+ and CD8+ T cells are elicited following infection and they are effective against secondary influenza virus infections (44, 45). There may be epitopes on the HA that could elicit broadly cross-reacting T cells that help clear virally infected cells (46, 47). In addition, influenza virus elicited T cells to internal gene products, such as NP of both the H1N1 and H3N2 viruses, could recognize epitopes on the internal genes of the Sw/MO/06 challenge virus. Further studies are needed to identify the exact mechanism of protection that these pre-immune animal models.

Across all of the pre-immune groups, the Z1 vaccinated ferrets had significantly higher cross-reactive H2 antibody titers compared to the other vaccination groups. The Z1 vaccinated ferrets had the highest average HAI titers and recognized more H2 strains than the other vaccines across all of the different pre-immunities. The Z1 vaccinated ferrets also had the highest average neutralization titers to more H2 influenza viruses regardless of the pre-immune background. The COBRA H2 HA vaccine is likely outperforming the wild-type H2 HA vaccines because they have more diverse epitopes. Higher diversity of epitopes in the COBRA HA would more likely elicit B-cells that cross-react across different antigenic sites on the H2 HA. Higher diversity of epitopes is beneficial for vaccinating people who are all pre-immune to either H1N1 and/or H3N2 influenza viruses. Vaccinating with a COBRA H2 HA antigen with highly diverse cross-reactive epitopes on a single antigen would increase the likelihood that multiple cross-reactive B cells will be retained in long-term immunological memory.

The Z1 COBRA HA also outperformed the Z5 COBRA and the two wild-type vaccines. Z1 outperforming Z5 was somewhat surprising since there are only four amino acids that differ between the two HA sequences. However, these four amino acids are spread across three of the seven antigenic sites on the H2 HA molecule (48, 49). It is likely that these mutations are altering the structure of multiple epitopes that are recognized by B cells and resulting in decreased antibody cross-reactive binding to other H2 HA proteins. Therefore, the Z1 COBRA HA would be an ideal vaccine candidate for a future stockpiled H2 influenza virus vaccine and administered to protect people from a future H2 influenza virus pandemic.

## Acknowledgments

The authors would like to thank Dr. Ivette A Nunez, Dr. Ying Huang, James D Allen, and Dr. Hyesun Jang for technical assistance. The authors thank the CVI protein production core for providing technical assistance in purifying recombinant HA proteins, as well as the University of Georgia Animal Resource staff, technicians, and veterinarians for excellent animal care.

## Author Contributions

Z. Beau Reneer - Conceptualization, Formal analysis, Methodology, Writing

Amanda L. Skarlupka - Formal analysis, Writing

Parker J. Jamieson – Formal analysis, Writing

Ted M. Ross – Conceptualization, Formal analysis, Funding acquisition, Methodology, Writing

## Supplemental Figures

**Supplemental Figure 1: Amino acid diversity in antigenic sites of WT and COBRA H2 HA sequences**

The amino acid differences for the WT and COBRA H2 HA sequences in the six H2 HA antigenic sites are shown in the six tables. Amino acids are numbered based upon H3 numbering system. Only amino acid positions with differences are shown. All of the other amino acids in the antigenic sites are the same for all of the H2 HA sequences used in this study.

**Supplemental Figure 2. Main effects of vaccine received, established pre-immunity and day post-infection on the log 10 viral nasal wash titer**

ANOVA (Outcome = Titer; Predictors = Vaccine + pre-immunity + Day) adjusted by Tukey Honest Significance Difference method for the effect sizes (ie difference between the means) for vaccines (A), pre-immunity (B), and day (C) when controlling for the main effects of the other variables. The horizontal line at 0.0 indicates identical means with no measured difference. If the 95% confidence intervals extended over this line the difference between the two compared groups are not significant at the p = 0.05 level. Comparisons are colored based on adjusted p-value. Significance groups were determined from the effect size plots for the vaccine received (D), pre-immunity (E), and day (F). Groups that share a letter are not significant compared to the other.

**Supplemental Figure 3. Change in HAI titer from prime to boost vaccination**

Columns are divided by vaccine group, and rows are divided by pre-immunity. Data points from Day 14 to Day 42 are paired based one ferrets change in HAI titer for a virus in the HAI panel. Significance between groups are analyzed in Supplemental Table 1.

**Supplemental Figure 4. Main effects of vaccine received, established pre-immunity and virus tested on the log 2 HAI titer on Day 42**

ANOVA (Outcome = Titer; Predictors = Vaccine + Pre-immunity + Virus) adjusted by Tukey Honest Significance Difference method for the effect sizes (ie difference between the means) for vaccines (A), pre-immunity (B), and virus (C) when controlling for the main effects of the other variables. The horizontal line at 0.0 indicates identical means with no measured difference. If the 95% confidence intervals extended over this line the difference between the two compared groups are not significant at the p = 0.05 level. Comparisons are colored based on adjusted p-value. Significance groups were determined from the effect size plots for the vaccine received (D), pre-immunity (E), and virus (F). Groups that share a letter are not significant compared to the other.

**Supplemental Figure 5. Main effects of vaccine received, established pre-immunity and virus tested on the log 2 neutralization titer with pooled sera collected on Day 42**ANOVA (Outcome = Titer; Predictors = Vaccine + Pre-immunity + Virus) adjusted by Tukey

Honest Significance Difference method for the effect sizes (ie difference between the means) for vaccines (A), pre-immunity (B), and virus (C) when controlling for the main effects of the other variables. The horizontal line at 0.0 indicates identical means with no measured difference. If the 95% confidence intervals extended over this line the difference between the two compared groups are not significant at the p = 0.05 level. Comparisons are colored based on adjusted p-value. Significance groups were determined from the effect size plots for the vaccine received (D), pre-immunity (E), and virus (F). Groups that share a letter are not significant compared to the other.

**Supplemental Figure 6: Establishment of Pre-immunity in Ferrets**

HAI titers for pre-immune ferrets against the strains that were used to establish their influenza virus pre-immunity. Serum from each ferret was obtained on day 60 post-infection and tested against the listed viruses for each pre-immune group.

**Supplemental Figure 7: Viral Nasal Wash Titers for H3N2-H1N1 and H1N1-H3N2 Pre-immune Groups**

Nasal washes were performed on D1, D3, D5 and D7 post-infection. The titers are recorded as log10 PFU/mL. The H3N2-H1N1 pre-immune ferrets are shown in panels A-D. The H1N1-H3N2 pre-immune ferrets are shown in panels E-H. The height of the bars shows the mean while the error bars represent mean standard error.

## Supplemental Tables

**Supplemental Table 1**

Paired t-test of log2 HAI titers measured after prime (Day 14) and boost (Day 42) vaccinations. The Holm correction was used to adjust for multiple comparisons. Samples were paired based on the ferret. T-tests were conducted to determine if HAI titers changed after stratified by pre-immunity and vaccine received. * p < 0.05; ** p < 0.01; *** p < 0.001

